# Striatal cell-type–specific molecular signatures reveal therapeutic targets in a model of dystonia

**DOI:** 10.1101/2024.10.07.617010

**Authors:** Kaitlyn M. Roman, Ashok R. Dinasarapu, Suraj Cherian, Xueliang Fan, Yuping Donsante, Nivetha Aravind, C. Savio Chan, H.A. Jinnah, Ellen J. Hess

## Abstract

Striatal dysfunction is implicated in many forms of dystonia, including idiopathic, inherited and iatrogenic dystonias. The striatum is comprised largely of GABAergic spiny projection neurons (SPNs) that are defined by their long-range efferents. Direct SPNs (dSPNs) project to the internal globus pallidus/substantia nigra reticulata whereas indirect pathway SPNs (iSPNs) project to the external pallidum; the concerted activity of both SPN subtypes modulates movement. Convergent results from genetic, imaging and physiological studies in patients suggest that abnormalities of both dSPNs and iSPNs contribute to the expression of dystonia, but the molecular adaptations underlying these abnormalities are not known. Here we provide a comprehensive analysis of SPN cell-type–specific molecular signatures in a model of DOPA-responsive dystonia (DRD mice), which is caused by gene defects that reduce dopamine neurotransmission, resulting in dystonia that is specifically associated with striatal dysfunction. Individually profiling the translatome of dSPNs and iSPNs using translating ribosome affinity purification with RNA-seq revealed hundreds of differentially translating mRNAs in each SPN subtype in DRD mice, yet there was little overlap between the dysregulated genes in dSPNs and iSPNs. Despite the paucity of shared adaptations, a disruption in glutamatergic signaling was predicted for both dSPNs and iSPNs. Indeed, we found that both AMPA and NMDA receptor-mediated currents were enhanced in dSPNs but diminished in iSPNs in DRD mice. The pattern of mRNA dysregulation was specific to dystonia as the adaptations in DRD mice were distinct from those in parkinsonian mice where the dopamine deficit occurs in adults, suggesting that the phenotypic outcome is dependent on both the timing of the dopaminergic deficit and the SPN-specific adaptions. We leveraged the unique molecular signatures of dSPNs and iSPNs in DRD mice to identify biochemical mechanisms that may be targets for therapeutics, including LRRK2 inhibition. Administration of the LRRK2 inhibitor MLi-2 ameliorated the dystonia in DRD mice suggesting a novel target for therapeutics and demonstrating that the delineation of cell-type–specific molecular signatures provides a powerful approach to revealing both CNS dysfunction and therapeutic targets in dystonia.

## Introduction

Dystonia is characterized by involuntary muscle contractions that cause debilitating twisting movements and postures.^1^ There are currently few effective treatments for dystonia because the underlying neuronal dysfunction is not completely understood. However, the basal ganglia are implicated in many forms of dystonia.^2–4^ A common cellular mechanism underlying basal ganglia dysfunction in dystonia is abnormal nigrostriatal dopamine neurotransmission. Pathogenic variants in genes critical for dopamine synthesis, vesicular packaging or reuptake in the presynaptic terminal cause dystonia.^5–10^ Correspondingly, pathogenic variants in genes critical for postsynaptic dopamine receptor signal transduction are associated with dystonia.^11–15^ Even without overt gene defects, disruptions in dopamine neurotransmission can cause dystonia, such as tardive dystonia following dopamine receptor antagonist treatment.^16^ Compelling evidence for the role of dopamine dysfunction in dystonia is DOPA-responsive dystonia (DRD), a group of childhood-onset dystonias that markedly improve after administration of L-DOPA, the synthetic precursor of dopamine. DRD is caused by pathogenic variants in genes necessary for catecholamine synthesis, including *GCH1* (GTP cyclohydrolase 1) and *TH* (tyrosine hydroxylase).^5, 6, 17, 18^ As a consequence, dopamine concentrations in DRD are significantly reduced,^6, 19, 20^ similar to the low dopamine levels in adult-onset Parkinson’s disease,^21^ yet the outcome is instead dystonia, suggesting that the specific motor dysfunction is not likely caused by the dopamine deficit per se. Instead, the motor outcome may be determined by the presynaptic dopaminergic deficit in combination with unique postsynaptic adaptations.

Dopamine plays a pivotal role in the control of movements by modulating the activity of the basal ganglia. The principal input structure of the basal ganglia is the striatum. Convergent dopaminergic and glutamatergic signaling in the striatum is integrated by GABAergic spiny projection neurons (SPNs). SPNs are broadly divided into two subtypes based on their long-range axonal projections and gene expression profiles. In general, direct SPNs (dSPNs) express D1 dopamine receptors (D1Rs) and project to the internal globus pallidus/substantia nigra reticulata (GPi/SNr) while indirect pathway SPNs (iSPNs) express D2 dopamine receptors (D2Rs) and project to the external pallidum (GPe).^22, 23^ Although dSPNs and iSPNs are largely segregated into distinct pathways, they act in concert to mediate and refine movements.^24–29^ Evidence suggests that both dSPNs and iSPNs are abnormal in dystonia. Functional imaging studies have shown that D1R and D2R signaling is abnormal in both inherited and idiopathic dystonias.^17, 30–35^ Further, recordings in patients demonstrate that long-range signaling in the axonal targets of dSPNs (GPi/SNr) and iSPNs (GPe) is abnormal.^36–41^ Because each SPN subtype is implicated in dystonia, a precise account of striatal cell-type–specific molecular adaptations is necessary to attain a comprehensive understanding of the mechanisms underlying dystonia.

A challenge to understanding cell-type–specific molecular changes in dystonia is the complexity of striatal anatomy. dSPNs and iSPNs are intermingled so whole tissue transcriptional profiling is not useful for identifying SPN subtype-selective changes underlying dystonia. Therefore, we used translating ribosome affinity purification (TRAP) to distinguish cell-type–specific molecular signatures in the translatome of genetically identified dSPNs or iSPNs in a mouse model of DRD. DRD knockin mice recapitulate the core features of the human disorder, including a human DRD-causing pathogenic variant in the *TH* gene (c.1141C > A),^7^ reduced tyrosine hydroxylase activity accompanied by a reduction in striatal dopamine concentrations and dystonic movements that improve in response to L-DOPA.^42^ Further, signaling defects in both SPN subtypes contribute to the expression of dystonia in DRD mice,^42^ consistent with findings in humans. Here, we demonstrate that SPNs exhibit cell-type–specific molecular adaptations in the DRD model of dystonia that are distinct from those that occur in a mouse model of Parkinson’s disease and demonstrate the utility of our findings for the identification of targeted therapeutics for the treatment of dystonia.

## Materials and methods

### Mice

For the DRD mouse TRAP studies, normal control (+/+) and homozygous DRD (*Th^DRD^*/*Th^DRD^*) littermates were used at 12 weeks of age, when the dystonia is most severe.^43^ DRD and control mice were F1 hybrids of C57BL/6J +/*Th^DRD^* x DBA/2J +/*Th^DRD^* to circumvent the high perinatal lethality exhibited by C57BL/6J DRD mice.^42^ The *Drd1a-EGFP/Rpl10a* transgenic line (RRID:IMSR_JAX:030254; Drd1a-TRAP) and the *Drd2-EGFP/Rpl10a* transgenic line (RRID:IMSR_JAX:030255; Drd2-TRAP) were used to isolate translating mRNA from dSPNs and iSPNs, respectively. The TRAP transgenes inbred on C57BL/6J were bred individually as hemizygotes onto the C57BL/6J +/*Th^DRD^* strain prior to the F1 hybrid cross to produce control and DRD experimental mice. For visualized *ex vivo* physiology, the transgenic BAC line *Drd1a-tdTomato* (RRID:IMSR_JAX:016204) was used to identify dSPNs^44^ and the transgenic BAC line *Drd2-eGFP* (RRID:MMRRC_000230-UNC) was used to identify iSPNs.^45^ *Drd1a-tdTomato* or *Drd2-eGFP* BAC transgenes inbred on C57BL/6J were crossed onto DRD mice as described for the TRAP transgenes. We have previously shown that neither *Drd1a-tdTomato* nor *Drd2-eGFP* transgene affects the dystonic movements observed in DRD mice.^46^ For the 6-hydroxydopamine (6-OHDA) experiment, mice were F1 hybrids of C57BL/6J x DBA/2J and hemizygous for the *Drd1a-EGFP/Rpl10a* transgene to replicate the background of DRD mice. Mice were bred at Emory University and maintained on a 12h light/dark cycle with *ad libitum* access to food and water and genotyped using PCR (see Supplementary Table 1 for primers). All experimental procedures were approved by the Animal Care and Use Committee at Emory University and Northwestern University and followed guidelines set forth in the *Guide for the Care and Use of Laboratory Animals*.

### 6-OHDA lesions

Sham-lesioned (n = 7 females, 8 males) and 6-OHDA-lesioned mice (n = 8 females, 8 males) were used for dSPN TRAP. Mice (6-7 weeks of age) were anesthetized with isoflurane (Piramal Healthcare, Mumbai, India), injected subcutaneously (s.c.) with meloxicam SR (4 mg/kg, ZooPharm) and mounted in a stereotaxic apparatus. A midline incision was made over the skull and a small hole was drilled over the right medial forebrain bundle. A Hamilton syringe with a 32-gauge needle was used to deliver a unilateral injection of 0.6 µL of 5 µg/µL 6-OHDA HBr in 0.02% ascorbic acid or vehicle over 5 minutes (from bregma: AP -1.2 mm; lat -1.1mm; DV -4.75mm). After infusion, the needle was left in place for an additional 5 minutes. The incision was closed with wound glue (Vetbond; 3M, St. Paul, MN). Four weeks post-surgery, the lesions were confirmed by scoring 5 min videos of spontaneous 360° rotations of mice placed in a cylinder. 6-OHDA-treated mice with ≥75% of rotations in the ipsilateral direction were considered hemiparkinsonian.

To determine the extent of the dopamine depletion typically induced using this surgical protocol, a separate group of mice was treated with 6-OHDA (n = 10 females, 11 males) and striatal dopamine concentrations were assessed by high pressure liquid chromatography in the Emory HPLC Bioanalytical Core using an ESA 5600A CoulArray equipped with an MD-150 × 3.2 mm C18, 3 µm column (Thermo Scientific) and a 6210 electrochemical cell (ESA, Bedford, MA) with potentials −175, 100, 350 and 425 mV. Dopamine was identified by matching retention time to known standards (Sigma Chemical Co., St. Louis MO) and quantified by comparing peak areas to those of the standard on the dominant sensor.

### Translating ribosome affinity purification (TRAP)

TRAP was performed as described.^47^ Sex-matched pairs of control and experimental mice were processed in parallel. Mice were euthanized by cervical dislocation, brains were rapidly extracted, and the striatum was dissected. For studies with DRD and control littermates, left and right striatum were collected and pooled from each individual mouse. For studies with hemiparkinsonian and sham-lesioned littermates, only the right (lesioned) striatum was collected. Translating RNA was isolated >6 weeks after unilateral sham- or 6-OHDA-lesion to ensure that we profiled chronic postsynaptic changes in gene expression, rather than neuroinflammation which lasts >30 days after lesion.^48, 49^ Immediately after dissection, striata were washed in ice-cold dissection buffer 1x HBSS (2.5 mM HEPES-KOH, pH 7.3, 35 mM glucose, 4 mM NaHCO_3_, and 100 µg/mL cycloheximide). Tissue from each mouse was individually homogenized in tissue-lysis buffer (20 mM HEPES-KOH, pH 7.3, 150 mM KCl, 10 mM MgCl_2_, 0.5 mM dithiothreitol, 100 µg/ml cycloheximide, EDTA-free Protease Inhibitor Mini-Complete (Roche), and 10 µl/ml SUPERase•In RNase Inhibitor (Ambion) using a glass teflon homogenizer at 900 RPM at 4°C. Nuclei and debris were removed with centrifugation at 2000 x g for 10 min at 4°C. 1,2-dihexanoyl-sn-glycero-3-phosphocholine (30 mM, Avanti Polar Lipids, Alabastar, AL) and NP-40 (1% vol/vol) were added to the supernatant and samples were incubated on ice for 10 minutes. The mixture was centrifuged at 20,000 x g for 10 minutes at 4°C. The supernatant was collected and resuspended in an affinity matrix containing anti-GFP monoclonal antibodies (HtzGFP-19F7 and HtzGFP-19C8; Antibody and Bioresource Core Facility, Memorial Sloan Kettering Cancer Center, New York, NY) bound to biotinylated Protein L (Thermo Scientific)-coated Streptavidin MyOne T1 Dynabeads (Invitrogen). Samples were incubated at 4°C overnight with gentle end-over-end mixing. After incubation, beads were washed four times with high salt buffer (20 mM HEPES-KOH, pH 7.3, 350 mM KCl, 10 mM MgCl2, 1% NP-40, 0.5 mM dithiothreitol, 100 µg/ml cycloheximide). RNA was extracted from mRNA-ribosome-bead complexes using the Absolutely RNA Nano Prep Kit (Agilent) according to the manufacturer’s protocol. Beads were vortexed in 100 µL of Nano Prep lysis buffer and 0.7 µL β-mercaptoethanol and then incubated for 10 min at room temperature. A magnet was used to separate the beads from the RNA in solution. Sulfolane (40% vol/vol) was added to the RNA and the mixture was transferred to a nano-spin cup where the RNA bound to a silica-based fiber matrix. The samples were treated with DNAase and washed to remove DNase and other proteins. RNA was eluted from the fiber matrix in 20 µL of elution buffer at 60°C. Samples were immediately frozen at -80°C and provided to the Molecular Evolution and High Throughput Sequencing Core at the Georgia Institute of Technology for library preparation and sequencing. Libraries were constructed using the NEBNext Single Cell/Low Input RNA Library Prep Kit for Illumina. After assessing the quality of the libraries (fragment sizes 150 - 1000bp), equimolar concentrations of indexed libraries were pooled. Samples from the DRD cohorts were run on the Illumina NovaSeq 6000 (2x150 bp; ∼25 million reads/sample). Samples from the parkinsonian cohort were run on the Illumina NovaSeq with the SP flow cell (2x150 bp; ∼25 million reads/sample). Each TRAP cohort was batch processed to reduce technical noise.

### Bioinformatic analysis

To clean the reads in the FASTQ files, BBduk (sourceforge.net/projects/bbmap/) and Trimmomatic^50^ were used to remove adapter sequences and low-quality bases, respectively. MultiQC^51^ was used to ensure sample quality before the RNA-seq reads were mapped against the mouse reference genome (GRCm38) using STAR,^52^ retaining only uniquely mapped reads. Differential expression analysis was performed with DESeq2, and genes with a Benjamini-Hochberg (BH) adjusted p value < 0.1 and fold change ≥ |1.25| were considered differentially expressed.^53^ Robust principal component analysis (rPCA) was used to detect outliers with the “rrcov” R package^54^ after the regularized log (rlog) transformation of the DESeq2 normalized counts. Volcano plots were generated from protein-coding genes with R package EnhancedVolcano (https://github.com/kevinblighe/EnhancedVolcano) or GraphPad Prism. Gene Set Enrichment Analysis (GSEA)^55, 56^ was run using the DESeq2 normalized gene counts, after removing low expressed genes with edgeR:filterByExpr^57^ and mapping the mouse Ensembl gene IDs to the *Homo sapiens* homolog gene symbol with biomaRt.^58^ Pathways were considered enriched with a nominal p value < 0.1 and the FDR<0.25, adjusted for gene set size and multiple hypothesis testing.

### Quantitative real-time PCR

Striata were collected from an independent cohort of DRD mice and control littermates (n = 6/genotype) for validation studies and from a cohort of C57BL/6J mice (n = 4) to assess expression of *Drd1* and *Drd2* (Supplementary Table 2) and stored at −80 °C until use. Total RNA was isolated using TRIzol (Invitrogen) homogenization and chloroform layer separation. The clear RNA layer was then treated with in-column DNAase (PureLink RNA Mini Kit, Ambion). RNA concentration was determined with a NanoDrop. cDNA was reverse-transcribed from 1000 ng of total RNA (High-Capacity cDNA Reverse Transcription Kit, ABI). The qPCR reaction mixture consisted of 10 μL SYBR Select Master Mix, 2 μL each 5 uM forward and reverse primers (Supplementary Table 1), 6 μL water, and 2 μL cDNA template. Samples were heated to 50 °C for 2 min, 95 °C for 10 min, and 40 cycles of 95 °C for 15 s and 60 °C for 1 min, followed by a melt curve stage consisting of 95 °C for 15 s, 60 °C for 1 min, 95 °C for 30 s, 60 °C for 15 s. Analyses were performed using the ΔΔC(t) method.^59^ Samples were normalized to *Gapdh* (see Supplementary Table 1 for primers).

### Western blotting

Striata were dissected from an independent cohort of control and DRD mice and homogenized in 50 mM Tris, 150 mM NaCl, 0.1% SDS, 0.5% Na deoxycholate, 1% Triton X-100, pH 8.0 plus PhosSTOP and cOmplete™, Mini, EDTA-free Protease Inhibitor Cocktail (Roche). Protein concentrations were determined using a BCA assay (Thermo Scientific). 50 μg protein/sample was denatured at 95 °C for 5 min in Laemmli buffer with 2.5% ß-mercaptoethanol and separated with a 4–20% Mini-PROTEAN (Bio-Rad) polyacrylamide gel in 25 mM Tris, 192 mM glycine and 0.1% SDS running buffer. After transfer of the proteins to a nitrocellulose membrane in 25 mM Tris, 192 mM glycine and 20% methanol, the blot was blocked with Intercept Blocking Buffer (LI-COR Biosciences) and subsequently incubated with primary antibodies against FOSB (rabbit monoclonal anti-FOSB, Cell Signaling Technology #2251, 1:150), TH (rabbit polyclonal anti-TH, Pel-Freez Biologicals #P40101–0, 1:1000) and β-actin (mouse monoclonal anti-β-actin, Cell Signaling Technology #3700, 1:1000) in blocking buffer overnight at 4 °C. Secondary antibodies (donkey anti-mouse IRDye 680RD and goat anti-rabbit IRDye 800CW, LI-COR Biosciences, 1:10,000) were incubated for 1 h at 4 °C in blocking buffer. The blot was imaged on a LI-COR Odyssey (700 channel intensity of 2.0, 800 channel intensity of 5.0) and analyzed using Image Studio Lite (Version 5.2). ΔFOSB was normalized to β-actin.

### Visualized *ex vivo* electrophysiology

Mice (8-14 weeks of age) were anesthetized with a ketamine-xylazine mixture and perfused transcardially with ice-cold aCSF containing the following (in mM): 125 NaCl, 2.5 KCl, 1.25 NaH_2_PO_4_, 2.0 CaCl_2_, 1.0 MgCl_2_, 25 NaHCO_3_, and 12.5 glucose, bubbled continuously with carbogen (95% O_2_ and 5% CO_2_). The brains were rapidly removed, glued to the stage of a vibrating microtome (Leica Instrument) and immersed in ice-cold aCSF. Parasagittal slices containing the striatum were cut at a thickness of 240 μm and transferred to a holding chamber where they were submerged in aCSF at 37 °C for 30 min and returned to room temperature for recording. Slices were then transferred to a small-volume (∼0.5 ml) Delrin recording chamber that was mounted on a fixed-stage, upright microscope (Olympus). Neurons were visualized using differential interference contrast optics (Olympus), illuminated at 735 nm (Thorlabs), and imaged with a 60× water-immersion objective (Olympus) and a CCD camera (QImaging). Genetically defined neurons were identified by somatic eGFP or tdTomato fluorescence examined under epifluorescence microscopy with a daylight (6,500 K) LED (Thorlabs) and appropriate filters (Semrock).

Recordings were made at room temperature (20–22 °C) with patch electrodes fabricated from capillary glass (Sutter Instrument) pulled on a Flaming-Brown puller (Sutter Instrument) and fire-polished with a microforge (Narishige) immediately before use. Pipette resistance was typically ∼3–5 MΩ. For current-clamp recordings, the internal solution consisted of the following (in mM): 135 KMeSO_4_, 10 Na_2_-phosphocreatine, 5 KCl, 5 EGTA, 5 HEPES, 2 Mg_2_ATP, 0.5 CaCl_2_, and 0.5 Na_3_GTP, with pH adjusted to 7.25–7.30 with KOH. For voltage-clamp recordings of excitatory postsynaptic currents (EPSCs), a low-chloride internal solution consisted of the following (in mM): 125 CsMeSO_3_, 10 Na_2_-phosphocreatine, 5 tetraethylammonium chloride, 5 QX-314 Cl, 5 HEPES-K, 5 EGTA-K, 2 Mg_2_ATP, 0.5 CaCl_2_, 0.5 Na_3_GTP, 0.2% (w/v) biocytin, with pH adjusted to 7.25–7.30 with CsOH. Stimulus generation and data acquisition were performed using an amplifier, digitizer, and pClamp (Molecular Devices). For current-clamp recordings, the amplifier bridge circuit was adjusted to compensate for electrode resistance and was subsequently monitored. The signals were filtered at 1 kHz and digitized at 10 kHz. Chemicals were from Sigma-Aldrich except KMeSO_4_ (ICN Biomedicals) and Na_2_-GTP (Roche).

Corticostriatal EPSCs were evoked with electrical stimulation using parallel bipolar tungsten electrodes (Frederick Haer) placed in the cortex. To measure AMPA and NMDA receptor-mediated currents, the holding potential of recorded neurons was alternated between − 80 and +40 mV. AMPA receptor-dependent currents were measured from the peak amplitude of EPSCs at –80 mV. NMDA receptor-dependent currents were measured at 50 ms after the EPSC peak at +40 mV.

SR95531 and QX314Cl (Tocris) were dissolved in water and frozen at –30 °C prior to use. Each compound was diluted to the appropriate concentrations by adding to the perfusate immediately before the experiment.

### Identification of potential therapeutic targets

The Connectivity Map (CMap)^60^ was used to identify mechanisms of action predicted to correct the DRD mouse striatal gene expression signature. The Touchstone dataset, which is a reference set of experimental results generated using the L1000 platform of the Library of Integrated Network-based Cellular Signaturesv(LINCS) ^61, 62^ was queried with the top 150 positively and negatively regulated genes from dSPN and iSPN datasets (300 genes/dataset). Results were based on mechanism of action using the set_type = ‘MOA_CLASS.’ Mechanisms of action were ranked based on similarity or dissimilarity compared to the DRD mouse gene expression signature, which is computed as a connectivity score. We identified mechanisms of action with negative connectivity scores of <-1.0 with the filters QC-PASSED and QUALITIY_PASSED, indicating that the mechanism of action was predicted to reverse the gene expression abnormalities. We further selected mechanisms of action with negative connectivity scores for both dSPNs and iSPNs. If mechanisms of action appeared more than once in the results, only the top scores were retained for presentation to avoid redundancy.

### Abnormal movement assessments

A behavioral inventory was used to define the type of abnormal movement, including tonic flexion (forelimbs, hindlimbs, trunk, head), tonic extension (forelimbs, hindlimbs, trunk, head), clonus (forelimbs and hindlimbs), and twisting (trunk, head), as described.^42^ Abnormal movements were scored for 30 s at 10-min intervals for 60 min. Behavioral scores were calculated by summing the scores from all scoring bins.

### Drug challenge

DRD mice were challenged with the LRRK2 inhibitor MLi-2 (Sigma-Aldrich) in a crossover design study with 5 days between tests. Mice were randomly assigned to receive either vehicle (25% solutol/75% saline) or MLi-2 (50 mg/kg, s.c., 10 ml/kg) on the first day of the test. Abnormal movements were assessed starting 15 min after vehicle or MLi-2 administration. Locomotor activity was recorded simultaneously using an automated photobeam apparatus equipped with 12 infrared beams in a 4 x 8 grid (San Diego Instruments). Beam breaks were recorded every 5 mins for the entire test session. Experimenters were blinded to treatment.

### Statistical analyses

Analyses of the RNA-seq data are described above. GraphPad Prism (version 10.2.3) was used for statistical analyses of behavior, western blot, qPCR, and electrophysiology. Parametric tests were used for most analyses using GraphPad Prism after determining that QQ plots (normality distribution plots) were nearly linear and approximated the line of identity. One-tailed *t* tests were used if there was an *a priori* prediction for the direction of the effect, otherwise two-tailed tests were used. Because the *Spred3* qPCR data were not normally distributed, we tested for outliers using the ROUT test (Q=5%), which identified a single outlier. Removing the outlier restored normality and parametric statistics were applied to the cleaned data. The locomotor activity data were not normally distributed due to one mouse that was extremely active. Because locomotor hyperactivity is sometimes observed in DRD mice, rather than excluding the mouse, a nonparametric Wilcoxon matched pairs signed rank test was used. Statistical results and sample sizes are reported in the figure legends.

### Data availability

The data that support the findings of this study are available from the corresponding author, upon reasonable request.

## Results

### TRAP transgenes do not affect abnormal movements in DRD mice

To determine if the Drd1a-TRAP or Drd2-TRAP transgenes, which were used to isolate cell-type–specific translating mRNA, altered the expression of dystonia in DRD mice, abnormal movements were assessed in control and DRD mice hemizygous for the Drd1a-TRAP or Drd2-TRAP transgenes or transgene-negative littermates. As expected, abnormal movements were not observed in transgene-negative control mice or in control mice carrying either TRAP transgene. The dystonic movements observed in DRD mice carrying either Drd1a-TRAP or Drd2-TRAP transgenes were comparable to those of DRD littermates without the transgenes (Supplementary Fig. 1) suggesting that the transgenes did not introduce obvious confounds.

### Adaptations in dSPN gene expression in DRD mice

To determine the molecular adaptations in the translatome of dSPNs, translating mRNA was extracted from control and DRD mice expressing the Drd1a-TRAP transgene and RNA-seq was performed on samples from each individual mouse. rPCA did not identify any outliers, so all samples were included in downstream analyses (Fig. 1A). To determine if the cell-type–specific RNA enrichment was effective, the levels of *Drd1* mRNA, which is a marker of dSPNs, were compared to those of *Drd2* mRNA, which is a marker for iSPNs. Translating mRNA extracted from Drd1a-TRAP striata was enriched for *Drd1* by >30-fold over *Drd2* mRNA in both control and DRD mice (Fig. 1B). By contrast, in mRNA extracted from whole striatum, *Drd1* mRNA is only ∼3-fold more abundant than *Drd2* mRNA (Supplementary Table 2). Of the 14646 protein-coding genes detected, 1383 were differentially expressed with 716 upregulated and 667 downregulated genes in dSPNs of DRD mice compared to controls (fold change > 1.25 and BH adjusted p value < 0.1; Fig. 1 C & D).

**Figure 1.**
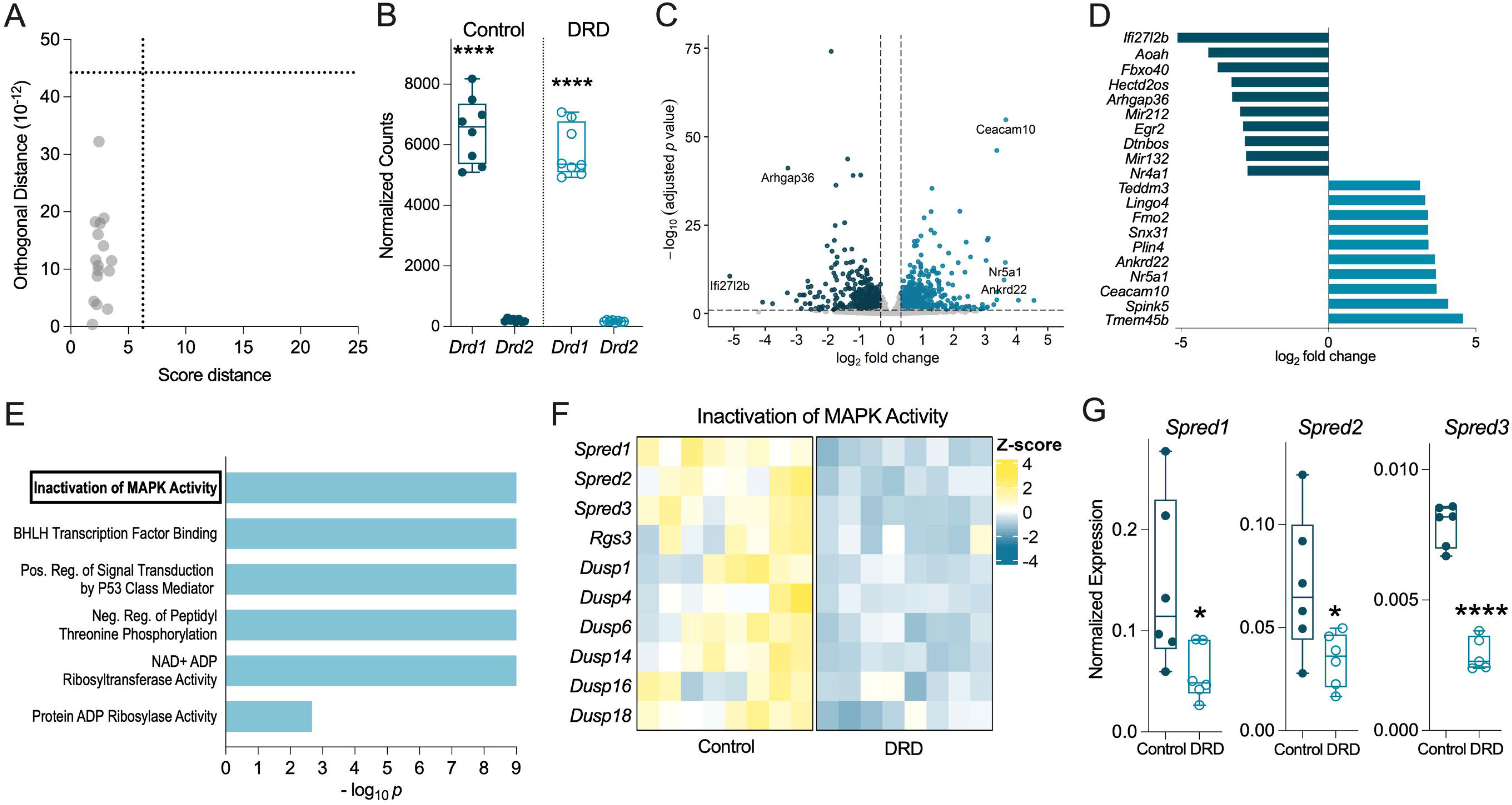
Adaptations in dSPN gene expression in DRD mice. **(A)** Robust principal component analysis outlier map. All samples (gray circles) were clustered within the cutoffs for score distance and orthogonal distance (represented by vertical and horizontal dotted lines, respectively) demonstrating no samples were outliers (n = 3 females, 5 males/genotype for panels A-F). **(B)** Assessment of TRAP enrichment for dSPN mRNA using normalized counts of *Drd1* and *Drd2* from RNA-seq on striatal mRNA extracted from control and DRD mice carrying the Drd1a-TRAP transgene. *Drd1* mRNA was significantly enriched compared to *Drd2* mRNA in both control (*t*_14_ = 16.22, *p* < 0.0001, one-tailed *t* test) and DRD mice (*t*_14_ = 18.40, *p* < 0.0001, one-tailed *t* test). **(C)** Volcano plot depicting protein-coding genes differentially expressed in dSPNs in DRD compared to control mice. Downregulated genes in DRD mice are represented by dark aqua dots and upregulated genes in DRD mice are represented by light aqua dots (fold change > |1.25|; BH adjusted *p* value < 0.1). All other genes are represented by light grey dots. **(D)** The top ten downregulated (dark aqua) and top ten upregulated (light aqua) genes in dSPNs in DRD mice. **(E)** Significantly enriched pathways in dSPNs identified by gene set enrichment analysis. All dSPN pathways were downregulated in DRD compared to control mice. The x axis represents - log_10_ *p* values with a maximum cutoff of 9. **(F)** Heatmap of individual variation of differentially expressed genes in dSPNs in the *Inactivation of MAPK Activity* pathway between control and DRD mice. Data are represented as z-scores, with yellow depicting higher expression and aqua depicting lower expression. Each column represents an individual mouse. **(G)** Validation of RNA-seq results from dSPNs using mRNA extracted from whole striatum. Normalized expression (ΔΔCT) of *Spred* mRNAs identified in ‘**F**’ was measured by qPCR in an independent cohort of control and DRD mice (n=6/genotype). *Spred1* (*t*_10_ = 2.414, *p* = 0.0182, one-tailed *t* test), *Spred2* (*t*_10_ = 2.428, *p* = 0.0178, one-tailed *t* test), and *Spred3* (*t*_9_ = 11.02, *p* < 0.0001, one-tailed *t* test) mRNAs were significantly downregulated in DRD compared to control mice. For box and whisker plots, horizontal lines illustrate the minimum, 25^th^ percentile, median, 75^th^ percentile, and maximum with values for individual mice indicated by circles. Asterisks indicate \**p* < 0.05 and \*\*\*\**p* < 0.0001.

To identify biological pathways associated with the hundreds of differentially expressed genes in DRD dSPNs, we performed gene set enrichment analysis (GSEA). Six pathways were identified (FDR<0.25); all were downregulated in DRD compared to controls (Fig. 1E). To validate our findings, we focused on *Inactivation of MAPK Activity* because MAPK signaling has been implicated in movement disorders, including in DRD mice (Fig. 1F).^63–67^ All three Sprouty-related genes (*Spred1*, *Spred2*, *Spred3*), which encode inhibitors of MAPK signaling,^68^ were downregulated in dSPNs of DRD mice suggesting a robust and consistent effect. Therefore, qPCR was used to quantify the mRNA expression of each *Spred* gene in an independent cohort of DRD and control mice that did not carry the TRAP transgenes. The expression of all *Spred* genes was significantly lower in DRD compared to controls (Fig. 1G), confirming the RNA-seq results.

### Adaptations in iSPN gene expression in DRD mice

To determine the molecular adaptations in the translatome of iSPNs in DRD mice, TRAP was performed using control and DRD mice expressing the Drd2-TRAP transgene. rPCA did not detect outliers (Fig. 2A). Both control and DRD samples were enriched in *Drd2* mRNA over *Drd1* mRNA by >28-fold (Fig. 2B), whereas *Drd1* mRNA is more abundant than *Drd2* mRNA in whole striatal mRNA (Supplementary Table 2). Of the 12653 protein-coding genes detected, 1898 were differentially expressed with 941 upregulated and 957 downregulated in iSPNs of DRD mice compared to controls (fold change > |1.25| and BH adjusted *p* value < 0.1; Fig. 2 C & D).

**Figure 2.**
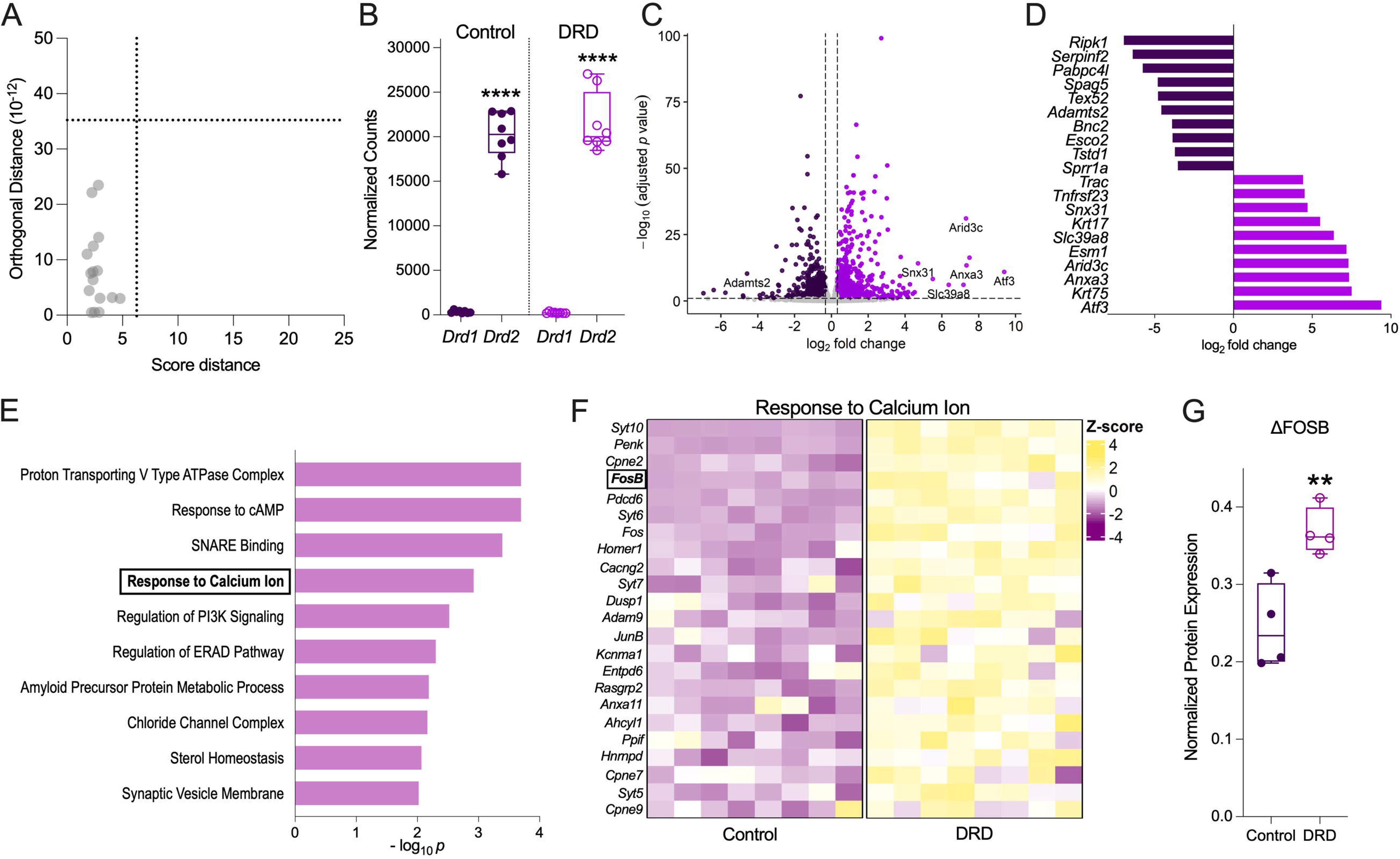
Adaptations in iSPN gene expression in DRD mice. **(A)** Robust principal component analysis outlier map. All samples (gray circles) were clustered within the cutoffs for score distance and orthogonal distance (represented by vertical and horizontal dotted lines, respectively) demonstrating no samples were outliers (n = 5 females, 3 males/genotype for panels A-F). **(B)** Assessment of TRAP enrichment for iSPN mRNA using normalized counts of *Drd2* and *Drd1* from RNA-seq on striatal mRNA extracted from control and DRD mice carrying the *Drd2*-TRAP transgene. *Drd2* mRNA was significantly enriched compared to *Drd1* mRNA in both control (*t*_14_ = 21.76, *p* < 0.0001, one-tailed *t* test) and DRD mice (*t*_14_ = 18.27, *p* < 0.0001, one-tailed *t* test). **(C)** Volcano plot depicting protein-coding genes differentially expressed in iSPNs in DRD compared to control mice. Genes significantly downregulated in DRD mice are represented by dark purple dots and genes significantly upregulated in DRD mice are represented by light purple dots (fold change > |1.25|; BH adjusted *p* value < 0.1). All other genes are represented by light grey dots. **(D)** The top ten downregulated (dark purple) and top ten upregulated (light purple) genes in iSPNs in DRD mice. **(E)** Significantly enriched, brain-relevant pathways in iSPNs identified by gene set enrichment analysis. All pathways identified for iSPNs were upregulated in DRD compared to control mice. **(F)** Heatmap of individual variation of differentially expressed genes in the *Response to Calcium Ion* pathway in iSPN-enriched mRNAs among control and DRD mice. Data are represented as z-scores, with yellow depicting higher expression and purple depicting lower expression. Each column represents an individual mouse. **(G)** Validation of RNA-seq results from iSPNs using western blot for ΔFOSB in striatal homogenates from control and DRD mice (n=4/genotype). Individual values of the densitometric quantification of ΔFOSB normalized to ß-actin for control and DRD mice are superimposed over box and whisker plots. ΔFOSB protein expression was significantly increased in DRD compared to control mice (*t*_6_ = 3.951, *p* = 0.0038, one-tailed *t* test). Box and whisker plots illustrate the minimum, 25^th^ percentile, median, 75^th^ percentile, and maximum with horizontal lines with values for individual mice indicated by circles. Asterisks indicate \*\**p* < 0.01 and \*\*\*\**p* < 0.0001.

All 112 GSEA pathways in iSPNs were upregulated in DRD mice compared to controls. This contrasts with dSPNs, where all biological pathways were downregulated. The top ten brain-relevant pathways included cellular processes such as intracellular signaling, and protein trafficking and degradation, among others (Fig. 2E). The *Response to Calcium Ion* pathway (Fig. 2F) was selected for validation because defects in calcium homeostasis are associated with many dystonias.^4^ Within this pathway, we focused on *Fosb* because it is implicated in other movement disorders.^69, 70^ *Fosb* encodes two alternatively spliced proteins, FOSB and ΔFOSB, which is the more stable isoform.^71^ Quantification of striatal ΔFOSB protein from an independent cohort of DRD and control mice using western blot analysis demonstrated that ΔFOSB protein was significantly increased in DRD compared to controls (Fig. 2G), in accordance with the differentially expressed *Fosb* mRNA.

### Cell-type–specific adaptations in dSPNs and iSPNs in DRD mice

Dopamine plays a critical role in the development and maturation of both SPN subtypes.^72, 73^ Therefore, the deficit in dopamine during development in DRD mice may result in similar adaptations in dSPNs and iSPNs. Of the upregulated genes in dSPNs (716 genes) and iSPNs (941 genes), only 106 mRNAs were upregulated in both SPN subtypes in DRD mice compared to controls (Fig. 3A). Like the upregulated genes, most downregulated genes in DRD mice differed between the SPN subtypes. Of the downregulated genes in dSPNs (667 genes) and iSPNs (957 genes), 88 genes were downregulated in both SPN subtypes in DRD mice (Fig. 3A). Notably, multiple differentially expressed genes in dSPNs and iSPNs were previously identified as differentially abundant in a limited proteomic analysis of bulk striatal homogenates in DRD mice compared to controls (Supplementary Table 3).^46^ Overall, few genes were regulated in parallel between the SPN subtypes.

**Figure 3.**
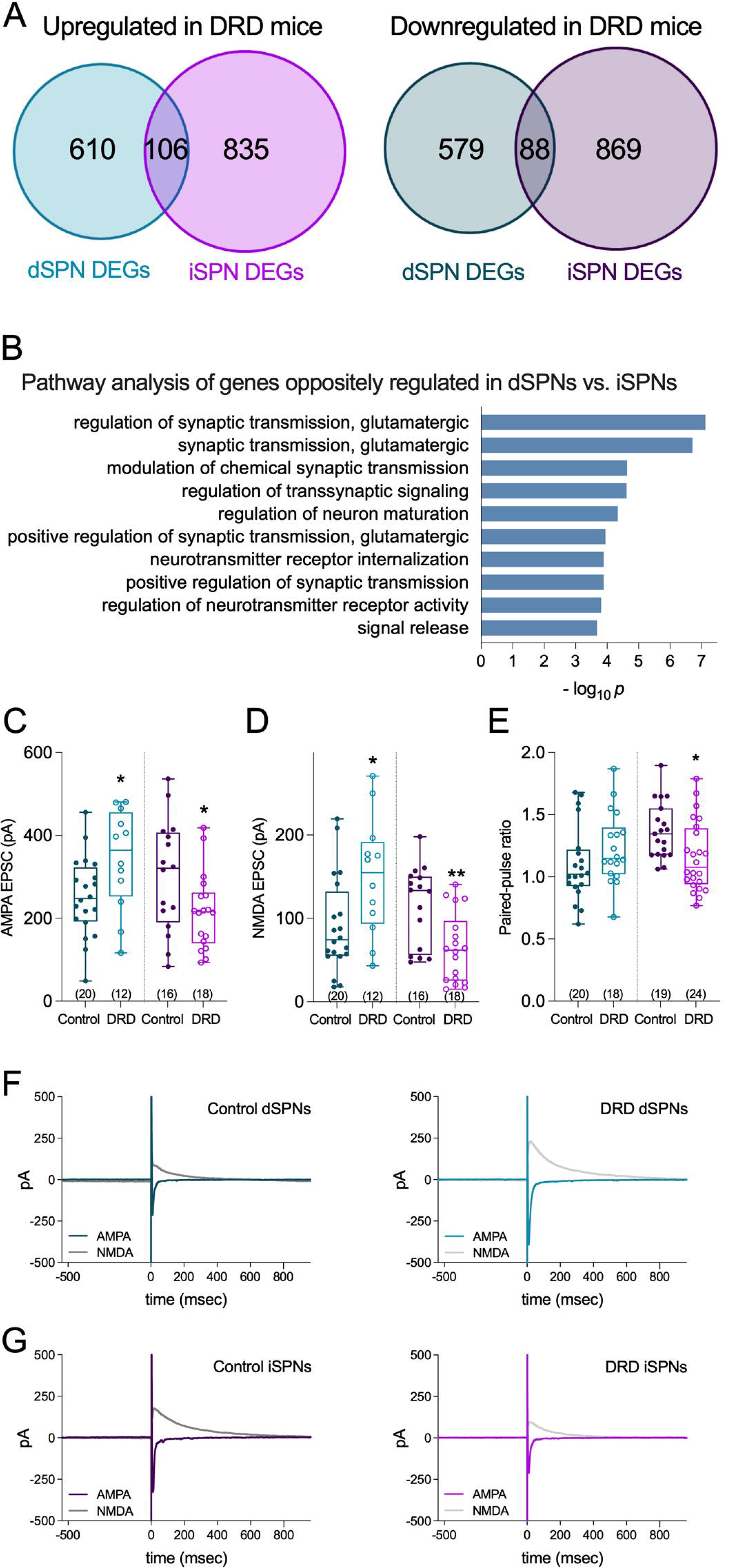
Cell-type–specific adaptations in dSPNs and iSPNs in DRD mice. **(A)** Venn diagram of dysregulated genes in dSPNs and iSPNs. 106 genes were upregulated in both dSPNs (light aqua) and iSPNs (light purple) while 88 genes were downregulated in both dSPNs (dark aqua) and iSPNs (dark purple) in DRD compared to control mice (fold change > |1.25|, BH adjusted p value <0.1). **(B)** Significantly enriched pathways identified by gene set enrichment analysis using genes that were regulated in opposite directions in dSPNs and iSPNs of DRD mice. **(C)** AMPA currents were increased in dSPNs (*t*_30_ = 2.346, *p* = 0.0256, two-tailed *t* test) but decreased in iSPNs (*t*_32_ = 2.179, *p* = 0.0368, two-tailed *t* test) of DRD mice compared to controls in response to electrically evoked EPSCs from corticostriatal synapses. **(D)** NMDA currents were increased in dSPNs (*t*_30_ = 2.560, *p* = 0.0158, two-tailed *t* test) but decreased in iSPNs (*t*_32_ = 3.303, *p* = 0.0024, two-tailed *t* test) of DRD mice compared to controls. **(E)** The paired pulse ratio (PPR) did not change in dSPNs (*t*_36_ = 1.339, *p* = 0.1888, two-tailed *t* test) but decreased in iSPNs (*t*_41_ = 2.623, *p* = 0.0122, two-tailed *t* test) of DRD mice compared to controls. For **C** and **D**, sample sizes appear in parenthesis. Box and whisker plots illustrate the minimum, 25^th^ percentile, median, 75^th^ percentile, and maximum with horizontal lines. Asterisks indicate \**p* < 0.05 and \*\**p* < 0.01 compared to control. **(F, G)** Representative traces for AMPA and NMDA components of EPSCs when SPNs were held at -80 and +40 mV, respectively, in dSPNs (F) and iSPNs (G) of control and DRD mice.

The dopaminergic deficit in DRD mice reduces signaling at both facilitatory G_s/olf_-coupled D1 dopamine receptors on dSPNs and inhibitory G_i_-coupled D2 dopamine receptors on iSPNs suggesting that translating mRNA may be oppositely regulated between these two cell types. The expression of 189 genes was oppositely regulated in dSPNs and iSPNs. A GSEA of these genes identified pathways associated with neurotransmission, with regulation of glutamatergic transmission accounting for the top two pathways (Fig. 3B), suggesting the hypothesis that glutamatergic neurotransmission in DRD mice is regulated in opposite directions in dSPNs compared to iSPNs. To test this hypothesis, we measured AMPA and NMDA receptor-mediated currents using visualized *ex vivo* physiology to record from genetically identified dSPNs or iSPNs from control and DRD mice. Using a standardized stimulation intensity to stimulate input from the sensorimotor cortex, we found that both AMPA and NMDA currents were increased in dSPNs of DRD mice (Figs. 3C, 3D, 3F), despite a decrease in excitability as reflected by the rheobase (Supplementary Fig. 2). On the contrary, in iSPNs, both AMPA and NMDA currents were decreased in DRD mice (Figs. 3C, 3D, 3G), consistent with a decrease in excitability (Supplementary Fig. 2). The changes in both the size of EPSCs and excitability in iSPNs may be postsynaptic adaptations to compensate for the increase in presynaptic release as suggested by the decreased paired-pulse ratio (Fig. 3E), which is a measure of the probability of transmitter release. Interestingly, both dSPNs and iSPNs appear to sustain a higher level of firing in DRD mice compared to controls.

### Molecular adaptations in dystonia vs parkinsonism

Although both DRD and Parkinson’s disease are caused by deficits in presynaptic dopamine concentrations, the presentation of these movement disorders is distinct. DRD is a childhood-onset hyperkinetic disorder, whereas Parkinson’s disease is an adult-onset hypokinetic disorder. Thus, a deficit in presynaptic dopamine per se does not entirely account for the clinical outcome. We, therefore, hypothesized that low presynaptic dopamine concentrations result in unique postsynaptic adaptations in DRD vs parkinsonism. To test this, we used the same TRAP approach that was used for DRD mice in the 6-OHDA mouse model of parkinsonism to provide a direct comparison between DRD and parkinsonism. We focused on dSPNs because immunohistochemical studies have demonstrated that ERK/MAPK signaling is dysregulated in dSPNs in both DRD mice^66^ and parkinsonian mice,^74–77^ which suggests the alternative hypothesis that postsynaptic adaptations do not distinguish between the disorders. Before sample collection, mice were tested for spontaneous circling behavior. All 6-OHDA-treated mice exhibited ≥75% of turns towards the lesioned hemisphere, confirming the dopamine lesions (Fig. 4A). Because postsynaptic adaptations are likely dependent on the extent of the presynaptic dopamine deficit, we also determined the efficacy of our lesion protocol by assessing striatal dopamine concentrations in a separate group of 6-OHDA-lesioned mice. Dopamine concentrations in the 6-OHDA-lesioned striata were <1% of unlesioned striata (Supplementary Table 4), which is comparable to DRD mice where striatal dopamine concentrations are also <1% of normal.^42^

**Figure 4.**
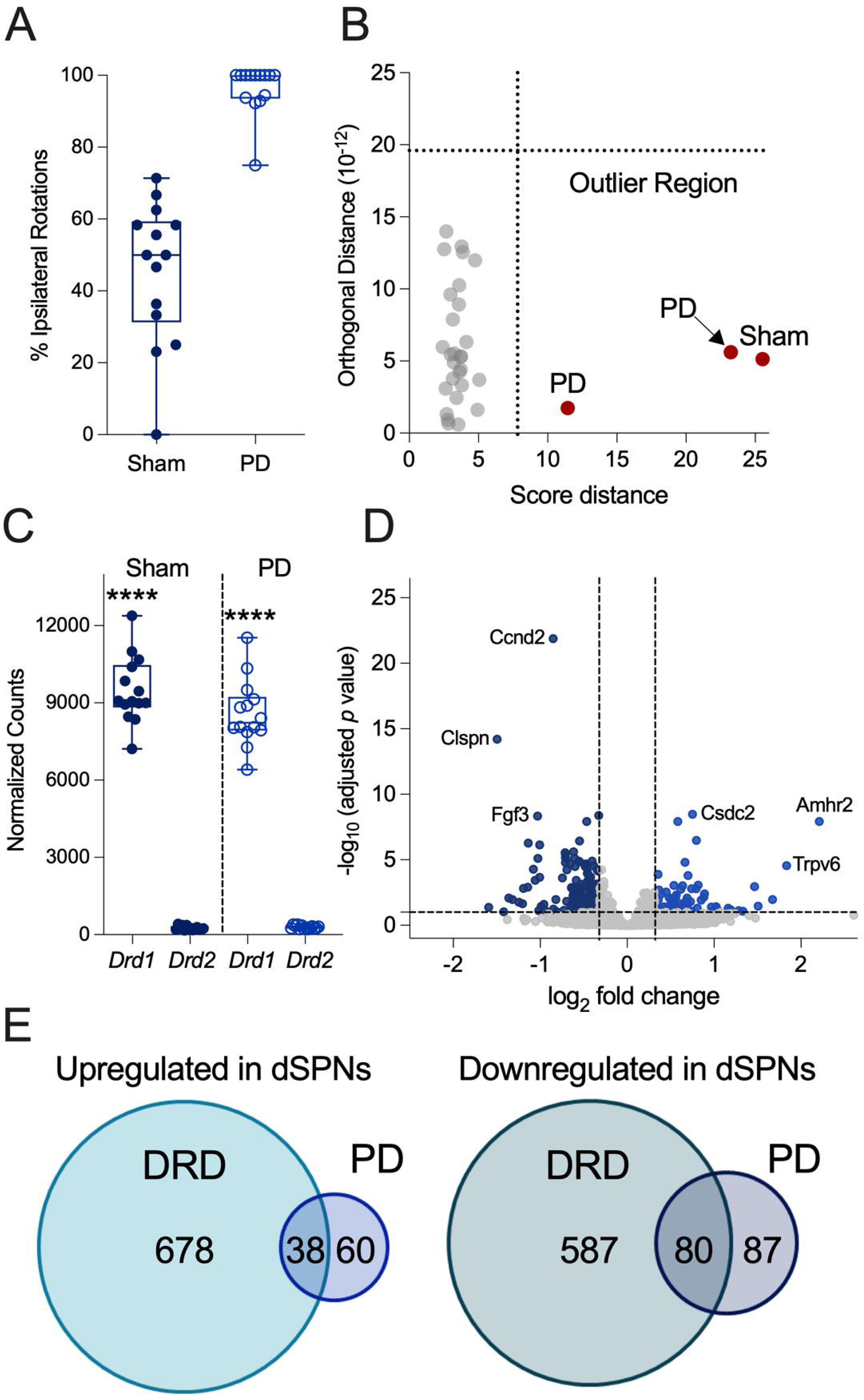
Molecular adaptations in dystonia vs parkinsonism. **(A)** Turning asymmetry in control (solid circles) and parkinsonian (PD; open circles) mice carrying the Drd1a-TRAP transgene for isolation of translating mRNA from dSPNs. Spontaneous 360° rotations were recorded over 5 minutes and ipsilateral rotations toward the 6-OHDA or sham-injected hemisphere were calculated as a percent of the total. **(B)** Robust principal component analysis outlier map. Most samples (gray circles) were clustered within the cutoffs for score distance and orthogonal distance (represented by vertical and horizontal dotted lines, respectively) while three samples (red circles) were outliers and excluded from subsequent analyses. **(C)** Assessment of TRAP enrichment for dSPN mRNA using normalized counts of *Drd1* and *Drd2* from RNA-seq on striatal mRNA extracted from control and 6-OHDA-lesioned mice carrying the Drd1a-TRAP transgene (n = 6 females, 8 males/treatment). *Drd1* mRNA was significantly enriched compared to *Drd2* mRNA in both sham-treated (*t*_26_ = 26.76, *p* < 0.0001, one-tailed *t* test) and 6-OHDA-treated mice (*t*_26_ = 24.22, *p* < 0.0001, one-tailed *t* test). **(D)** Volcano plot depicting protein-coding genes differentially expressed in dSPNs in 6-OHDA-treated compared to control mice. Genes significantly downregulated in 6-OHDA-treated mice are represented by dark blue dots and genes significantly upregulated in 6-OHDA-treated mice are represented by light blue dots (fold change > |1.25|; BH adjusted *p* value < 0.1). All other protein-coding genes are represented by light grey dots. **(E)** Venn diagrams of upregulated and downregulated genes in dSPNs in DRD and parkinsonian (PD) mice. Thirty-eight genes were upregulated in both conditions and 80 genes were downregulated in both conditions. Box and whisker plots illustrate the minimum, 25^th^ percentile, median, 75^th^ percentile, and maximum with horizontal lines with values for individual mice indicated by circles. Asterisks indicate \*\*\*\**p* < 0.0001.

Translating RNA from dSPNs in sham and 6-OHDA lesion samples (n = 14 mice/condition after removal of 3 outliers detected by rPCA; Fig. 4B) was enriched in *Drd1* mRNA compared to *Drd2* mRNA, demonstrating that the TRAP procedure was effective (Fig. 4C). Of the 12782 protein coding genes detected, 265 were differentially expressed in 6OHDA-lesioned compared to sham-lesioned striata (fold change >1.25 and BH adjusted p value < 0.1; Fig. 4D), which is comparable to another translatomic study of dSPNs in 6-OHDA-lesioned mice,^65^ but markedly less than the 1383 dysregulated genes in dSPNs of DRD mice. A comparison of the differentially expressed genes revealed that of the 716 upregulated genes in DRD dSPNs and the 98 upregulated genes in 6-OHDA-lesioned dSPNs, only 38 genes were upregulated in dSPNs in both conditions (Fig. 4E). Of the 667 downregulated genes in DRD mice and the 167 downregulated after 6-OHDA lesion, only 80 genes were downregulated in dSPNs in both conditions (Fig. 4E). Notably, a GSEA of the shared genes identified ‘MAPK cascade’ as the top biological pathway (padj = 0.077), consistent with previous immunohistochemical studies.^66, 74–77^ These results suggest that while some adaptations are shared, the majority are unique to each disorder.

### Identification of potential therapeutic targets for dystonia

Treatments for the dystonias are largely unsatisfactory. Therefore, to identify potential therapeutic targets, differentially expressed genes from either dSPNs or iSPNs in DRD mice were used as an input to query CMap using its CLUE tool (https://clue.io), which contains gene expression profiles from cell lines exposed to either small molecule compounds or genetic manipulations.^60^ For this chemical genomics approach, we probed the database to identify compounds expected to normalize gene expression (indicated by a negative connectivity score^55^) in both dSPNs and iSPNs because the translatome of each cell type was dysregulated in DRD mice. The drug mechanisms with the lowest connectivity scores in both dSPNs and iSPNs were inhibitors of enzyme or receptor activity (Fig. 5A). Although several intracellular signaling molecules were identified, no single signaling cascade was clearly implicated and most proteins lack cell-type specificity. However, the identification of ‘LRRK (leucine-rich repeat kinase) inhibitor’ as a mechanism for attenuating abnormalities in DRD mice was notable because the expression of the Parkinson’s disease-associated gene *LRRK2* is relatively specific to the striatum. Indeed, *LRRK2* is abundantly expressed in both dSPNs and iSPNs and mediates dopamine receptor intracellular signaling.^78, 79^ We found that the expression of *Lrrk2* is dysregulated in both dSPNs and iSPNs in DRD mice whereby *Lrrk2* was significantly downregulated in dSPNs but upregulated in iSPNs (Fig. 5B). Therefore, we tested the centrally acting selective LRRK2 inhibitor MLi-2 for antidystonic effects in DRD mice. Administration of 50 mg/kg MLi-2, a dose that is known to inhibit LRRK2 activity in mouse brain,^80^ significantly reduced dystonic movements in DRD mice (Fig. 5C). Importantly, MLi-2 did not affect locomotor activity suggesting that the dose was well-tolerated and did not induce competing behaviors such as akinesia or sleep (Fig. 5C).

**Figure 5.**
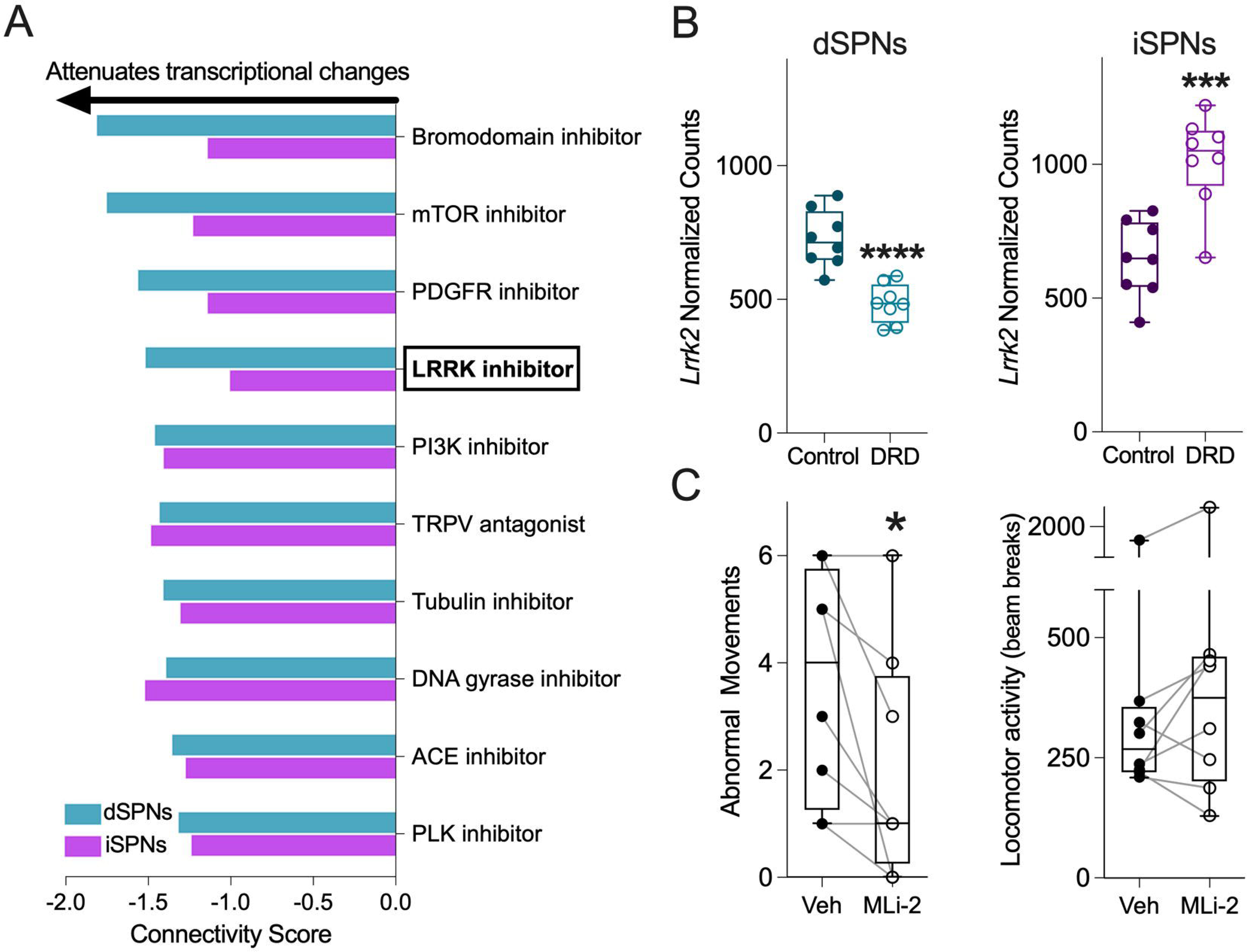
Identification of potential therapeutic targets for dystonia. **(A)** Mechanisms of action predicted to correct both dSPN and iSPN gene expression in DRD mice were identified using CMAP. Negative connectivity scores of <-1.0 are predicted to reverse the gene expression abnormalities. **(B)** Normalized expression of *Lrrk2* translating mRNA in dSPNs and iSPNs of control and DRD mice (n = 8/genotype for each cell type). *Lrrk2* expression was significantly reduced in dSPNs of DRD mice compared to control mice (padj < 0.0001) but significantly increased in iSPNs of DRD mice compared to control mice (padj = 0.00027). **(C)** The LRRK2 inhibitor MLi-2 attenuates dystonia in DRD mice. Abnormal movements and locomotor activity were assessed simultaneously after administration of vehicle or MLi-2 in a crossover design (n = 5 females, 3 males). Compared to vehicle, MLi-2 significantly reduced abnormal movements in DRD mice (*t*_7_ = 2.728, *p* = 0.0147, paired one-tailed *t* test). Compared to vehicle, MLi-2 did not affect locomotor activity (*p* = 0.3125, two-tailed Wilcoxon test). Data from individual mice are plotted in open circles with lines indicating the response to vehicle and compound. Box and whisker plots illustrate the minimum, 25^th^ percentile, median, 75^th^ percentile, and maximum with horizontal lines with values for individual mice indicated by circles. \**p* < 0.05, \*\*\**p* < 0.001, \*\*\*\**p* < 0. 0001.

## Discussion

Here, we identified the adaptations in direct and indirect pathway SPNs translatomes in a mouse model of DRD, identifying over 1300 dysregulated genes with distinct adaptations in each neuronal subtype, providing a molecular signature of the striatal dysfunction in a model of DRD. In addition to dystonia, striatal dysfunction underlies several movement disorders including Parkinson’s disease, Huntington’s disease and L-DOPA-induced dyskinesias. Studies using TRAP to isolate the translatome of dSPNs and iSPNs from mouse models have revealed distinct adaptations in each disorder. In a previous study using the 6-OHDA model of Parkinson’s disease, Heiman *et al*. identified <250 dysregulated genes in the translatomes of dSPNs and iSPNs,^75^ similar to our findings in dSPNs in the 6-OHDA model. By contrast, in a model of early stage Huntington’s disease, >5000 genes were dysregulated in each SPN subtype.^81^ Although the number of differentially expressed genes is comparable between dSPNs and iSPNs in models of parkinsonism, Huntington’s disease and DRD, responses are imbalanced in a model of L-DOPA-induced dyskinesias with >4500 dysregulated genes in dSPNs but <450 dysregulated genes in iSPNs.^65^ The unique molecular signatures of dSPNs and iSPNs in each movement disorder provides an opportunity to dissect the cellular features that give rise to the distinguishing features of these disorders and to identify disease-specific therapeutic targets.

In dSPNs of DRD mice, the MAPK pathway was disinhibited. This is in line with other research investigating the impact of abnormal dopaminergic neurotransmission. Altered ERK/MAPK signaling is observed in dopamine-deficient mice,^82^ and in dSPNs from parkinsonian mice.^65^ Consistent with disinhibition of MAPK signaling, supersensitivity of ERK phosphorylation in response to dopaminergic agonist challenge has been observed in DRD mice,^66^ suggesting a common mechanism by which dSPNs adapt to deficits in dopamine neurotransmission. In iSPNs, dysregulated pathways included protein trafficking and degradation, transmembrane ion transport, translation initiation, and mitochondrial function, among others. Notably, the endoplasmic reticulum (ER)-associated degradation (ERAD) pathway was dysregulated. The ERAD pathway translocates misfolded proteins to the cytosol for degradation by the proteasome. Dysregulation of the ERAD pathway is hypothesized to increase the sensitivity of neurons to ER stress, and several dystonia-associated-genes converge on pathways associated with the ER stress response.^83–87^ Dysregulation of genes associated with calcium handling was also identified in iSPNs in DRD mice. Abnormal calcium handling was first proposed as a mechanism underlying dystonia based on observations that pharmacologically- or genetically-mediated disruptions in calcium channel activity induce generalized dystonia in mice.^88, 89^ Subsequently, pathogenic variants in multiple genes involved in calcium homeostasis were identified in individuals with dystonia, including *ANO3*, *CACNA1A*, *CACNA1B*, *CAMK4*, *CAMTA1*, *HPCA*, *KCNMA1*, and *KCTD17.*^4, 90–97^

Our results suggest that glutamatergic neurotransmission is enhanced in dSPNs but diminished in iSPNs in DRD mice, a premise suggested by gene expression data and supported by electrophysiologic experiments. Ensembles of dSPNs and iSPNs receive spatially and temporally organized glutamatergic afferents from the cortex. This clustering of excitatory activity is thought to be critical for encoding and selecting movements.^24, 25, 98–100^ Impaired action selection is a hallmark of dystonia whereby attempted voluntary movements exacerbate the involuntary abnormal movements and extraneous unintentional movements often co-occur with dystonic movements in unaffected body regions (motor overflow). In DRD mice, an imbalance in the glutamatergic responses in dSPNs and iSPNs may not only disrupt long-range signaling at axonal targets (i.e., GPe, GPi/SNr), but may also degrade the spatiotemporal organization of intrastriatal microcircuits, resulting in aberrant action selection.

Both DRD and parkinsonism are characterized by a deficit in presynaptic dopamine yet the molecular signatures of dSPNs were unique to each disorder. While reduced dopamine neurotransmission occurs in both dystonia and Parkinson’s disease, the striking differences in postsynaptic adaptations might be explained by the differences in developmental period in which this perturbation occurs. Neural development in DRD mice occurs in the presence of < 1% of normal dopamine concentrations due to the mutation in *Th* whereas in hemiparkinsonian mice, the loss of dopamine neurotransmission does not occur until basal ganglia circuitry is mature.

Dopamine plays a critical role in embryonic development by providing trophic support to nascent striatal neurons.^72^ Later, in the early postnatal period, striatal dopaminergic signaling mediates the development of SPN intrinsic excitability,^73^ which is abnormally reduced in both SPN subtypes in DRD mice (Supplementary Fig. 2). By contrast, in parkinsonism, the intrinsic excitability of dSPNs is increased while the intrinsic excitability of iSPNs is reduced.^101^ Thus, the expression of parkinsonism versus dystonia likely results from dopamine depletion in combination with very specific and unique adaptions in SPNs that may be determined by the timing of the dopaminergic deficit.

Despite the strong association between dopamine neurotransmission and dystonia, dopaminergic drugs are not generally used to treat dystonia, in large part due to the heterogeneity in the etiology of the dystonias and thus the unpredictable response among patients.^102^ The identification of LRRK2 as a potential therapeutic target suggests an alternative strategy for the treatment of dystonias. While *Lrrk2* mRNA is expressed at low levels in the substantia nigra, it is unlikely that MLi-2 exerted its effects by altering presynaptic dopamine transmission in DRD mice considering that dopamine concentrations are <1% of normal. In contrast, *Lrrk2* is abundantly expressed in postsynaptic SPNs,^79, 103–105^ where it negatively regulates PKA activity through a direct interaction with the regulatory subunit PKARIIβ.^78^ LRRK2 inhibition or *Lrrk2* knockout activates PKA^106^ and facilitates D1R signaling in dSPNs.^107^ Reducing the negative regulation of PKA with a LRRK2 inhibitor may alleviate dystonia by enhancing D1R intracellular signaling, which is attenuated in DRD mice due to the mutation-induced dopamine deficit. Indeed, we have previously demonstrated that potentiating D1R signaling through D1-like receptor agonist treatment ameliorates the dystonia in DRD mice.^42^ While most dystonias are not associated with a reduction in brain dopamine concentrations like that observed in DRD, many dystonias are associated with abnormal dopamine neurotransmission arising from a variety factors that cannot be remediated by L-DOPA, such as abnormal release or dopamine receptor dysregulation. That targeting LRRK2, an enzyme downstream of dopamine receptor signal transduction, was effective in a dystonia caused by a presynaptic defect in dopamine synthesis suggests that therapeutics designed to circumvent the various upstream instigating defects could be effective in many forms of dystonia.

This study has several limitations. Gross anatomical differences between mutant and control mice could confound the results, but we have previously demonstrated that both the size of the striatum and the density of dSPNs and iSPNs are comparable between DRD and control mice.^46^ Although mRNA abundance is not always predictive of protein expression, ribosome-attached translating mRNA, as was used here, is a better reflection of protein expression than total mRNA.^108^ Next, the abundance of some mRNAs varies across subregions of the striatum,^109–115^ but it was not possible to resolve regional variation in differentially expressed genes using TRAP. It is also important to note that D2Rs are expressed in cholinergic interneurons as well as iSPNs. However, because only ∼2% of striatal neurons are cholinergic interneurons,^116^ this is unlikely to impact the results.

This work is focused on striatal dysfunction, which is implicated in many forms of dystonia. Current models suggest that dystonia is a network disorder involving basal ganglia, cerebellum, thalamus, cortex, and other motor regions. Dystonia may arise from dysfunction in one or more brain regions or the interaction between regions.^2, 117–120^ While the molecular signatures will vary with the locus of dysfunction, here, we have demonstrated that the delineation of cell-type–specific molecular signatures provides a powerful approach to revealing both CNS dysfunction and therapeutic targets in dystonia.

## Acknowledgements

The Emory University School of Medicine HPLC Bioanalytical Core (EHBC) provided technical support for the quantitation of striatal dopamine concentrations.

