## Supplementary material for "Striatal cell-type–specific molecular signatures reveal therapeutic targets in a model of dystonia"

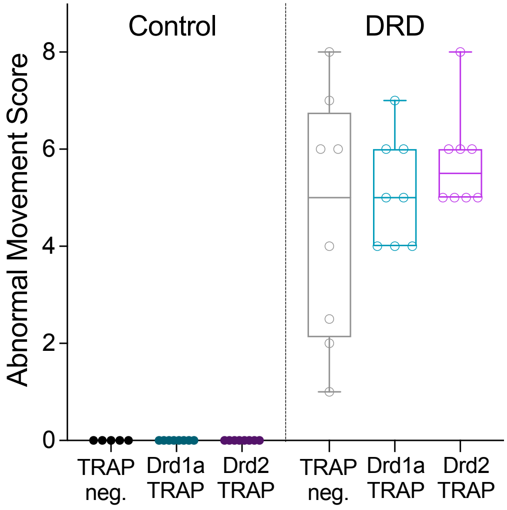


**Figure S1. TRAP transgenes do not affect the expression of abnormal movements in DRD mice.** Abnormal movements were assessed in control and DRD mice in the presence or absence of either the Drd1a-TRAP or Drd2-TRAP transgene (n = 5-8/genotype). Experimenters were blinded to transgene genotype. Neither transgene affected abnormal movements in DRD mice (One-way ANOVA, $F_{2, 21}$ = 0.9547, $p = 0.4010$). Control mice did not exhibit abnormal movements regardless of transgene status. Mice depicted here were susequently used to for cell-type specific translating mRNA analyses. Box and whisker plots illustrate the minimum, 25$^{th}$ percentile, median, 75$^{th}$ percentile, and maximum with horizontal lines. Values for individual mice are indicated by circles.

Figure S2

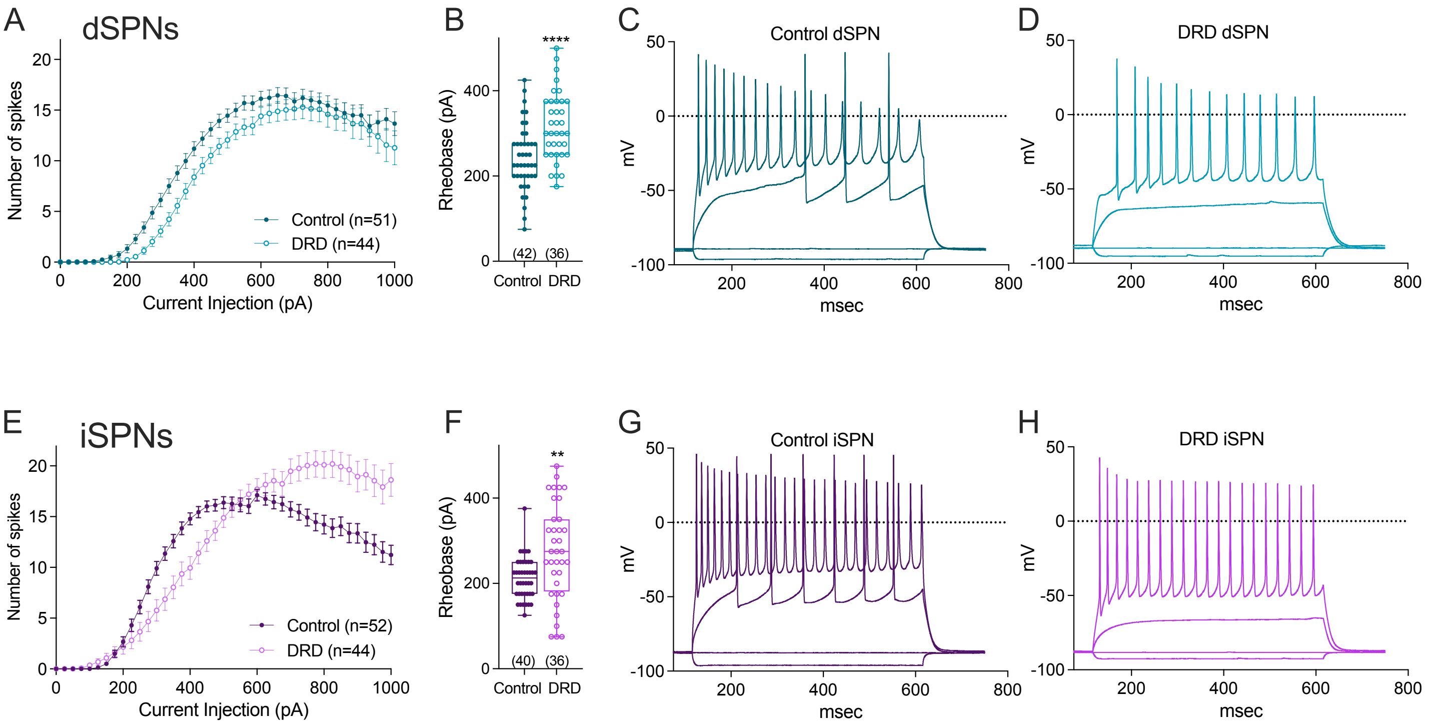


**Figure S2**. Excitability of dSPN and iSPNs is reduced in DRD mice. Input-output relationships (f-I curve) of dSPNs **(A)** and iSPNs **(E)** demonstrating a shift to the right and reduction in excitability in DRD mouse SPNs compared to controls. Rheobase was significantly increased in dSPNs **(B)** and iSPNs **(F)** in DRD mouse SPNs compared to controls (dSPNs: $t_{76} = 4.225$, $p < 0.0001$, two-tailed *t* test ; iSPNs: $t_{74} = 30211$, $p = 0.0020$, two-tailed *t* test). Horizontal lines illustrate the minimum, 25$^{th}$ percentile, median, 75$^{th}$ percentile, and maximum with values for individual cells indicated by circles. Numbers in partheses indicate sample size. $**p < 0.01$ and $****p < 0.0001$. **(C, D, G, H)** Representative current clamp traces in response to current injections of -150, 0, 250 and 625 pA.

### Supplementary Tables

#### Supplementary Table 1 PCR primer sequences

| Gene | Forward Primer | Reverse Primer |
|---|---|---|
| *Th*$^{DRD}$ | ACACCGAAGCAGAGACTGT | CTGATGCTACTTCTCCAGG |
| *EGFP/Rpl10a* | GCACGACTTCTTCAAGTCCGCCATGCC | GCGGATCTTGAAGTTCACCTTGATGCC |
| *Drd1a-tdTomato* | GTGGGCTCTATGGCTTCTG | CACGCTGATCACACAGAGG |
| *Drd2-eGFP* | GAGGAAGCATGCCTTGAAAA | TGGTGCAGATGAACTTCAGG |
| *Drd1* | ATCGTCACTTACACCAGTATCTACAGGA | GTGGTCTGGCAGTTCTTGGC |
| *Drd2* | TGGCTGCCCTTCTTCATCACGC | TGAAGGCCTTGCGGAACTCAATGT |
| *Gapdh* | CATGTTTGTGATGGGTGTGA | TGCATTGCTGACAATCTTGA |
| *Spred1* | GAAGACACTTCCCGTTCCCTA | GCCTTGCTGACTGAATGGTAT |
| *Spred2* | CCGACGTTTCATCATTGGAAGG | CCCCTGTCAAAGGCTCGTG |
| *Spred3* | GGGGCACTACGTCATCCAC | GAACCTCGACTCAGGGCAG |

**Supplementary Table 2 Expression of *Drd1a* and *Drd2* mRNA in striatum**

| Gene | Normalized Expression |
|---|---|
| *Drd1a* | 0.06489 ± 0.001472488 |
| *Drd2* | 0.02193 ± 0.003315312 |

qPCR was performed using mRNA extracted from whole striatal homogenates from normal mice. Expression of mRNAs (ΔΔCT) was normalized to *Gapdh* (n = 4). Values represent mean ± SEM.

**Supplementary Table 3 Proteomic validation of differentially expressed genes in dSPNs or iSPNs in DRD mice.**

| Gene | Protein | dSPN mRNA | iSPN mRNA | Striatal Protein |
|---|---|---|---|---|
| *Actn2* | Actinin alpha 2 | ↑ | - | ↑ |
| *Dbn1* | Drebrin 1 | - | ↓ | ↓ |
| *Dgkb* | Diacylglycerol kinase, beta | ↓ | - | ↓ |
| *Hsd17b10* | Hydroxysteroid (17-beta) dehydrogenase 10 | ↑ | - | ↑ |
| *Hspa12a* | Heat shock protein 12A | - | ↓ | ↓ |
| *Lamp5* | Lysosomal-associated membrane protein family, member 5 | ↑ | ↑ | ↑ |
| *Mccc1* | Methylcrotonoyl-Coenzyme A carboxylase 1 (alpha) | ↑ | - | ↑ |
| *Me2* | Malic enzyme 2, NAD(+)-dependent, mitochondrial | - | ↑ | ↑ |
| *Ociad2* | OCIA domain containing 2 | - | ↑ | ↑ |
| *Prkcg* | Protein kinase C, gamma | ↓ | ↓ | ↓ |
| *Rem2* | Rad and gem related GTP binding protein 2 | - | ↑ | ↑ |
| *Sema4a* | Semaphorin 4A | - | ↑ | ↑ |
| *Sept6* | Septin 6 | - | ↓ | ↓ |
| *Spata2l* | Spermatogenesis associated 2-like | - | ↓ | ↓ |
| *Srm* | Spermidine synthase | - | ↑ | ↑ |
| *Tomm70a* | Translocase of outer mitochondrial membrane 70A | ↑ | ↑ | ↑ |

Striatal proteomic results are from Briscione *et al*[46] whereby 1805 proteins were identified from bulk striatal homogenates and 57 proteins were differentially expressed between control and DRD mice. Symbols indicated significantly upregulated (↑), significantly downregulated (↓), or no significant change (-).

**Supplementary Table 4 Striatal dopamine concentrations after unilateral 6-OHDA lesion in mice**

| | N | Intact side | Lesioned side (%) |
|---|---|---|---|
| Females | 10 | 121 ± 11 | 0.7 ± 0.1 (0.58%) |
| Males | 11 | 123 ± 8 | 0.5 ± 0.1 (0.40%) |

Dopamine concentrations (ng/mg protein) were assessed in males and females to ensure that the lesion was not sex-biased due to sex differences in body size. (%) indicates residual dopamine after lesion compared to the intact side. Data represent means ± SEM.
